# A Continuous-Time Semi-Markov Model for Animal Movement in a Dynamic Environment

**DOI:** 10.1101/353516

**Authors:** Devin S. Johnson, Noel A. Pelland, Jeremy T. Sterling

**Affiliations:** Alaska Fisheries Science Center, NOAA Fisheries, Seattle, Washington, U.S.A.

**Keywords:** Animal movement, semi-Markov model, dynamic habitat, model selection, multiple imputation, posterior model probability, resource selection

## Abstract

We consider an extension to discrete-space continuous-time models animal movement that have previously be presented in the literature. The extension from a continuous-time Markov formulation to a continuous-time *semi*-Markov formulation allows for the inclusion of temporally dynamic habitat conditions as well as temporally changing movement responses by animals to that environment. We show that with only a little additional consideration, the Poisson likelihood approximation for the Markov version can still be used within the multiple imputation framework commonly employed for analysis of telemetry data. In addition, we consider a Bayesian model selection methodology with the imputation framework. The model selection method uses a Laplace approximation to the posterior model probability to provide a computationally feasible approach. The full methodology is then used to analyze movements of 15 northern fur seal (*Callorhinus ursinus*) pups with respect to surface winds, geostrophic currents, and sea surface temperature. The highest posterior model probabilities belonged to those models containing only winds and current, SST did not seem to be a significant factor for modeling their movement.

## 1 Introduction

Studying the movement of animals has become ubiquitous in the ecological literature in the past decade. One traditionally thorny issue for developing statistical models for telemetry data is relating spatial habitat conditions to movement processes. This difficulty arises because of the mismatch in support for the telemetry data (continuous spatial domain) and the observed habitat variables (usually a spatial grid or raster) and the often random times when location is observed. As the habitat variables are the lowest spatial resolution of the two types, thus, it makes sense to bring the telemetry data to the level of the habitat data, so-to-speak. We propose a generalization of the spatially-discrete, continuous-time movement of Hanks et al. (2015) that can relate movement between cells to a dynamically changing environment evaluated at the cell level.

In recent years, there have been several methodologies based on translating telemetry data to a discrete version and modeling movement between cells based on temporally static habitat variables (see Hooten et al. 2010, Hanks et al. 2015, Hanks and Hughes 2016, and Avgar et al. 2016). In addition, Johnson et al. (2013) illustrated that for static environments spatial point process models could also be used by collapsing over the temporal index of the telemetry data.

The continuous-time Markov chain (CTMC) movement model of Hanks et al. (2015) is especially appealing due to the fact that the continuous time formulation means that the model interpretation is not subject to the choice of time discretization. In addition, the CTMC model can be fit with GLM or GAM software for efficient parameter estimation. The CTMC model has been used to analyze mountain lion (*Puma concolor*) and northern fur seal movements (*Callorhinus ursinus*) (e.g., Buderman et al. 2018; Hanks and Hughes 2016; Hanks et al. 2015). However, there some limitations in the CTMC model that prohibit use of temporally indexed covariates. We propose a generalization of the CTMC to a continuous-time *semi*-Markov chain (CTSMC) movement model for which habitat can vary in time.

The inspiration for this research is the migratory movements of northern fur seal pups in relation to space and time-dependent physical environmental conditions, namely, ocean surface currents, marine surface winds, and sea surface temperature (SST). Each of these habitat variables are derived from satellite remote sensing or atmospheric reanalysis products. These variables were chosen based on previous evidence of their influence on migratory movements or physiological processes in this or other similar pinniped species. Ream et al. (2005) found evidence in satellite-tagged adult female northern fur seals of migratory movement alignment with mesoscale oceanographic surface currents. Anecdotal evidence and traditional knowledge of the influence of marine winds on the migratory trajectories of northern fur seal pups extends back millennia, as documented in testimonials of Aleut hunters to US government officials during study of the Bering Sea northern fur seal population in the late 1800s (Hooper, 1895). These hunters were in agreement that seals always traveled with a fair wind and sea, that the prevailing patterns of these variables influenced the passes through which migratory animals exited the Bering Sea, and that the timing of fall storms could influence the onset of migration. Widespread satellite tagging has provided the opportunity to confirm and quantify these effects; Lea et al. (2009) found differing departure and dispersal patterns of Pribilof Islands northern fur seal pups in years of contrasting atmospheric forcing, and evidence of more rapid movement in the presence of a tailwind. Sterling et al. (2014) and Pelland et al. (2014) found statistical evidence that surface winds influence adult northern fur seals during migration, with high winds associated with linear, directed movement, though directionality in relation to the wind was not explored.

While no direct evidence of the influence of SST on migratory movement exists for this species, results from the study of metabolic rates of captive northern fur seal pups imply that given the climatological SST patterns in the North Pacific, their thermoregulatory capabilities are insufficient for them to remain in the Bering Sea and northwestern Gulf of Alaska in winter (Rosen and Trites, 2014). Water temperature has also been found to influence the metabolic rate and likely the fasting duration of newly-weaned Antarctic fur seal pups (Rutishauser et al., 2004), which have a similar morphology to northern fur seals and also undergo a long post-weaning migration through subarctic waters that initiates in the fall. The metabolic constraints imposed by cold SSTs, in combination with the observed general southward movement of northern fur seal pups toward warmer SST after their departure from the Pribilof Islands, suggest that SST is possibly an additional migratory environmental cue for this species.

In this paper our objectives are to develop the CTSMC model from first principles such that we do not need to rely on the distributional assumptions of Hanks et al. (2015) with respect to cell residence time (Section 2). Although, see Hooten et al. (2017, Sec. 7.4) for an alternative development of the CTMC model. Because the habitat can change before the animal moves cells we can not rely on a constant rate of movement while the animal remains in a cell. Following CTSMC development, in Section 3, we show that with only slight modification, the imputation and GLM approach of Hanks et al. (2015) can still be used for parameter inference. In addition to model fitting we illustrate methodology for Bayesian model selection using this 2 stage method. In Section 4, we illustrate the full methodology for fitting and model selection using migration data from 15 northern fur seal pups tagged in Alaska, USA.

## 2 Continuous-Time Semi-Markov Model for Movement

The description of the data and notation generally follows that of Hanks et al. (2015), however, we have made a few alterations to accommodate the temporal dynamics of the semi-Markov generalization of their model. In a continuous-time, discrete-space movement model the geographical study domain is partitioned into a cells indexed by 𝒢 = {1,…, *n*}. These cells can be thought of as a mathematical graph with nodes represented by the cell centroids. Each node, *i*, has a set of neighboring nodes 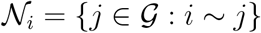, where *j* ~ *i* means that cells *i* and *j* are directly connected and the animal can move from cell *i* to cell *j* in one move. For example, neighboring cells might share a border in a raster partition, but, cells might also represent spatially separate patches that the animal might visit. As with all semi-Markov models, the complete path of the animal, 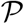, can decomposed into the times at which cell transitions are made (jump times), ***τ*** = {*τ*_0_, *τ*_1_, …, *τ*_*K*_}, and the sequence of cells visited (embedded Markov chain) **G** = {*G*_0_, …, *G*_*K*_}. We will assume that *τ*_0_ = 0 and *G*_0_ is the known starting cell. We use the notation *G*(*t*) to represent the location of the animal for any time in [0,*T*], e.g., *G*(*t*) = *G*_*k*_ for *t* ∈ [*τ*_*k*_, *τ*_*k*+1_).

### 2.1 General likelihood formulation

The heart of the CTSMC model is the emigration rate function

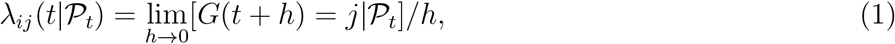

where 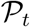 is the path of the animal up to time *t*, i.e., 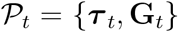, where *τ*_*t*_ = {*τ*_*k*_: *τ*_*k*_ < *t*} and **G**_*t*_ = {*G*_*k*_: *τ*_*k*_ < *t*}. Herein, we use the notation ‘[A|B]’ to represent the probability density (distribution) function of *A* given *B*. Therefore, 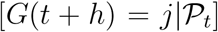 is the probability that *G*(*t* + *h*) = *j* given the path up to time *t*. The total rate of emigration from cell *i* at time *t* is the accumulation of the emigration rate 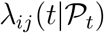 over 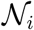,

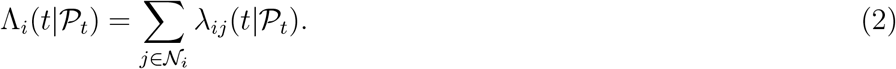

Using the total rate of emigration from a cell *i* at time *t*, we can obtain the distribution of the time of the next move, *τ*_*k*_ given the animal moved to cell i at time *τ*_*k*−1_. Using standard results from temporal point process methodology (see Hooten et al., 2017, Section 3.1.6), where the emigration rate function is mathematically equivalent to a point process intensity function, the density function of next movement time is,

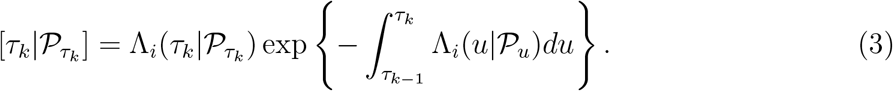

There is one complication that needs to be addressed at the end of the telemetry device deployment, the censoring of the last observation. If the individual is followed from time *τ*_0_ = 0 to time *T* ≥ *τ*_*K*_, the last cell movement, then the time between *τ*_*K*_ and *T* also needs to be accommodated. So, let *τ*_*K*+1_ be the unobserved ‘next’ transition occurring after the animal is no longer being observed at time *T*. The quantity necessary to accommodate the censoring is,

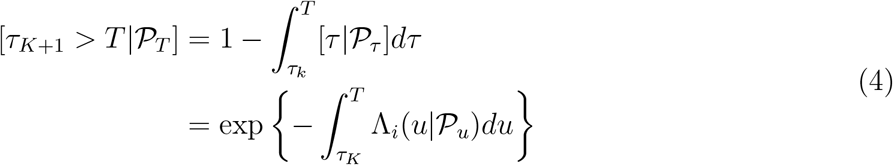

(Hooten et al., 2017). For the remainder of the model development we will notationally set *τ*_*k*+1_ = *T*, the end of the observation window for the telemetry deployment.

Now, if one conditions on the fact that the next move will be at time *τ*_*k*_, then the cell to which the move is made, is a categorical variable over 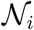,

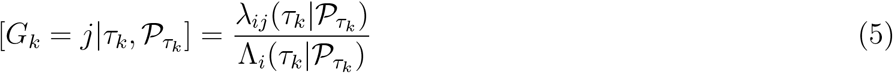

(Norris, 1998). If we now assume that the emigration hazard function, 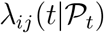, is also a function of a parameter vector, say *θ* then, given an observed path 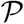, the likelihood is,

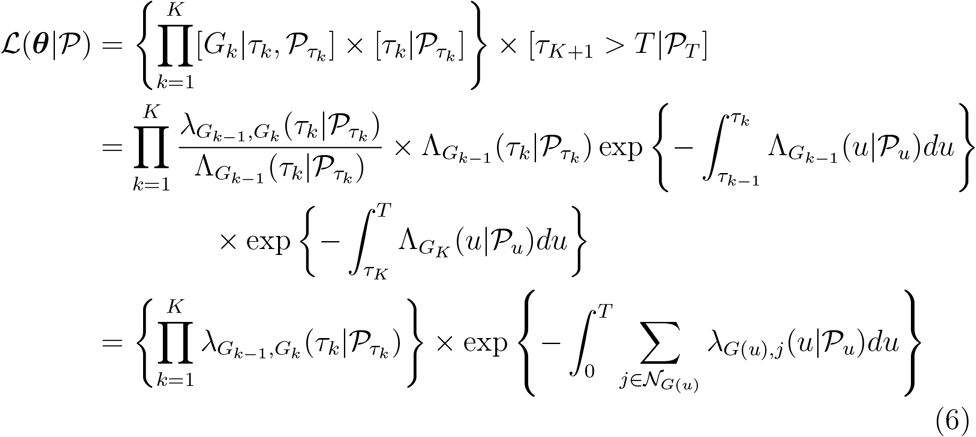

There are a couple of notes here,

### 2.2 A proportional rate model

Now that we have a general form for the likelihood of the movement process, we can consider specifying a model that includes the external variables which might be thought to influence movement. Here we follow the same model form given by Hanks et al. (2015) and Hanks and Hughes (2016), however, we generalize it to accommodate a time-varying structure and place it in a survival analysis context with a rich history of model development. In the survival modeling context, one of the most popular class is the proportional hazards models (Cox and Oakes, 1984). In the CTSMC movement context, for *G*(*t*) = *i* and 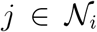, the proportional rates (hazards) model with a time-varying covariate is given by

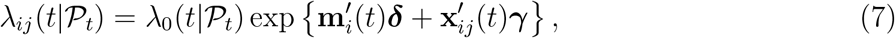

where 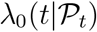 a a baseline hazard function, **m**_*i*_(*t*) is a covariate associated only with cell *i* at time *t*, **x**_*ij*_(*t*) is a covariate associated with both cell *i* and *j* (e.g., a gradient of an environmental variable) at time *t*, and ***δ*** and γ are regression coefficients associated with the respective covariates. The **m** variables are motility drivers in that they only affect the rate at which the animal will leave a cell, they have no influence on whether an animal is attracted to another cell; they disappear in equation (5). Whereas, the ‘gradient’ variables, **x**, influence rate of movement by drawing animals to other neighboring cells.

At this point, the model (7) is similar to that given by Hanks and Hughes (2016), however, there are some notable differences. First, and most obvious, the covariates are potentially time-varying. Second, the baseline transition rate need not be constant over time. The transition hazard can change with time since the last transition. This leads to clustered (or regular) transition times with respect to those that are exponentially distributed in the CTMC model. Another difference between the CTMC and CTSMC models are how the model treats temporally varying coefficients. In (7), the coefficients, ***δ*** and γ are constant though time. But, as with the Markov version, one might consider γ(*t*) or ***δ***(*t*). However, the difference in the CTSMC extension is that the coefficients can change smoothly through time even as the animal remains within the cell. The coefficients of the CTMC can only change when the animal transitions cells. This seems more mathematically appropriate, however, the realized difference in effect may be small in practice.

In addition to the obvious link between the CTSMC model and its CTMC special case, there is also a similarity with the continuous-space and timepoint process analysis of movement data presented by Johnson et al. (2013). The spatio-temporal point process likelihood is very similar to the CTSMC likelihood in (6) with the exception that the summation in the right hand side is replaced with an integration over the continuous spatial domain of neighboring sites. It is obvious that if we allow the cells to shrink in area while simultaneously increasing the number of neighbors to maintain equal neighborhood area, the summation will become an integral in the limit. Therefore, we can think of the CTSMC model as a spatially discrete approximation of the spatio-temporal point process of Johnson et al. (2013). The benefit of the CTSMC model is that by defining transitions between a small number of neighboring cells, the computation of the likelihood becomes much less demanding.

## 3 Statistical Inference

### 3.1 Likelihood calculation

The log-likelihood function in equation (6) is often challenging to evaluate in general. Hanks et al. (2015) present an approximation that allows users to obtain maximum likelihood estimates using standard GLM software for the CTMC model. Further, this approximation can be made as accurate as desired. In the context of survival analysis, this same approach was used by Holford (1980) and Laird and Olivier (1981). Following the same development used in the survival analysis, we can form an approximation for calculating the CTSMC movement model likelihood.

To begin the approximation, we first select a set of quadrature points, *q*_*i*_, to numerically evaluate the integral in (6). These quadrature points include times when cell transitions occurred, ***τ*** = {*τ*_0_,…,*τ*_*k*_}, and times where dynamic covariates changed values. In addition, the quadrature set also includes any additional points that the practitioner feels necessary to approximate the integral. For example, in Section 4, a 15 minute grid of times was also included in the quadrature set to model seal movement. Then, the approximation assumes 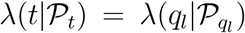 for *t* ∈ (*q*_*l*−1_, *q*_*l*_]. To ease the notational burden, we also set,

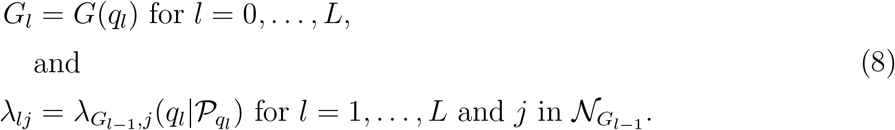

Now the integral is approximated with the summation,

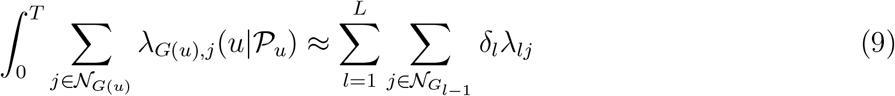

as arbitrarily close as desired, where *δ*_*l*_ = *q*_*l*_ − *q*_*l*−1_. After placing the approximation (9) into (5), one obtains the approximate likelihood function,

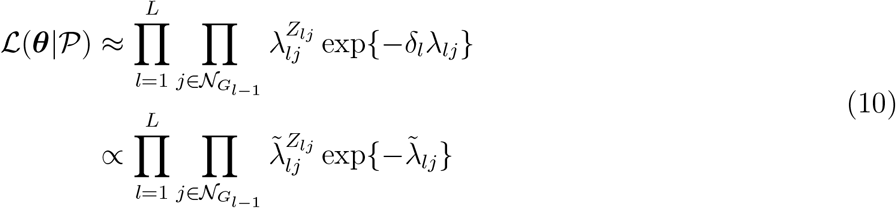

where 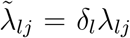 and *Z*_*lj*_ = 1 if *q*_*l*_ ∈ ***τ*** and *G*_*l*_ = *j* for some *k* = 1,…, *K*, that is, *q*_*l*_ is a transition time and *j* is the cell transitioned to at time *q*_*l*_, else *Z*_*lj*_ = 0. Notice, as with the model specification of Hanks and Hughes (2016) and Hanks et al. (2015), that the likelihood function formed by (10) is proportional to a Poisson likelihood function where *Z*_*lj*_ are the ‘data’ and 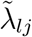 are the rate parameters. The only difference between this formulation and that of Hanks et al. (2015) is that there are times, *q*_*l*_ where *Z*_*lj*_ = 0 for all *j*, that is, there was no movement to a cell, but the λ_*ij*_ have changed.

### 3.2 GLM/GAM and the proportional rate model

Using the the Poisson approximation (10) one can gain powerful computational assistance if the proportional hazards model (7) is used. Because of the linear structure on the log scale, standard GLM software can be used to fit the CTSMC movement model. On the log scale, the Poisson rate used for the *Z*_*lj*_ data is

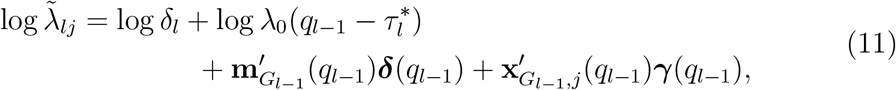

where log *δ*_*l*_ takes the form of a known offset and 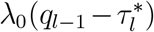 is the baseline cell transition rate that depends on 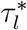 the time of the last transition prior to *q*_*l*_. Note, that we have now allowed the coefficients to vary with time now. Assuming λ_0_ is constant (we will discuss this momentarily), the parameters are all linear on the log scale, so any GLM fitting software can be used. One may also estimate time-varying coefficients using a ‘varying coefficients’ model as illustrated in the present analysis of northern fur seal migration in Section 4.

We now examine the baseline rate function, λ_0_. To begin, if one models log 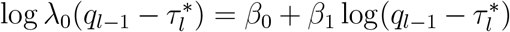 then the proportional rate model is still in a log-linear form, so standard GLM software can be used. This log-linear model for λ_0_ implies the waiting times between cell transitions follows a Weibull distribution if covariates are not considered (Cox and Oakes, 1984). If *β*_1_ < 0 then the rate function decreases with time since last transition and the times of transition will tend to be clustered in time (Hooten et al., 2017). The reverse is true for *β*_1_ > 0 transitions will occur at more regular intervals. For *β*_1_ = 0 the residence times will be exponentially distributed as in the CTMC model (Hanks and Hughes, 2016). If 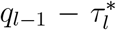 is used instead of the log version, the residence times will be distributed according to the Gompertz-Makeham distribution (Cox and Oakes, 1984). However, if neither of these simple models are useful, it is straightforward to model λ_0_ nonparametrically using a GAM model to directly estimate λ_0_ treating 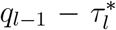 as a covariate to smooth.

### 3.3 Path uncertainty and the process imputation approach

Until this point, we have constructed the model and likelihood as if, in practice, we would observe 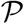 as data. This may be the case in the future as satellite telemetry devices improve, e.g., see Liu et al. (2014) for path reconstruction at subsecond intervals using accelerometer tags. However, for the most part, locations are observed sparsely and irregularly throughout the course of deployment. Therefore, we recommend the use of an *imputation* approach (Scharf et al., 2017) for analysis of traditional telemetry data in this framework. The imputation approach in movement analysis was initially proposed by Hooten et al. (2010) and is also the used by Hanks et al. (2015) for the Markov version of the discrete space movement model. Although not necessarily proposed for fully Bayesian inference, we take that approach here because it provides a coherent method of analysis from estimation to model selection.

The process imputation approach proceeds by considering the true quasi-continuous (in space and time) path of the animal, ***μ***.(t). If ***μ***(*t*) were observed for all *t* (***μ*** from here on) then we could summarize it into the data we desire, namely 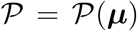. However, locations are usually only observed continuously in space, but sporadically in time with error, say, 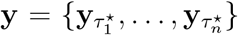, where *τ*\* is an observation time in [0, *T*]. To obtain the correct Bayesian posterior distribution for the movement parameters, ***θ***, the posterior distribution must be marginalized over the missing ***μ*** process, that is,

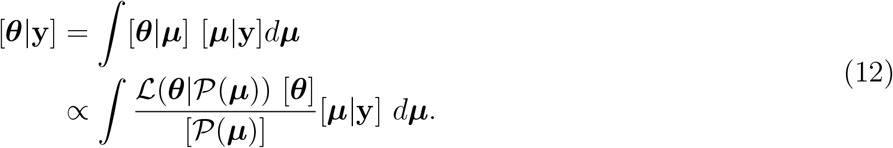

In practice, a two step procedure is used to describe [***θ***|**y**] by first drawing a sample from the imputation distribution ***μ*** ~ [***μ***|**y**], *r* = 1, …, *R*, computing the discrete space path from the imputed data, 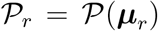, then summarizing 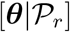 in some way. For example, Hanks et al. (2015) describes obtaining the posterior mean and variance of 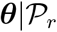 using the asymptotic distribution of the MLE, 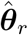 as an approximation of the posterior from which one can obtain the unconditional mean and variance of 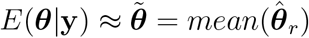 and 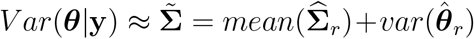, where 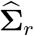 is the estimated asymptotic covariance matrix for the MLE 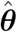. Herein we use ‘*mean*’ and ‘*var*’ to represent sample mean and variance over the imputation index, *r*.

As with imputation approaches in general, calculating interval estimates is not always straightforward (Rubin and Schenker, 1986), as there are many ways one may combine the data to obtain appropriate intervals. For example, Hanks et al. (2015) uses 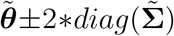 to obtain marginal intervals. However, this approach assumes that [***θ***|**y**] is generally normal looking to obtain 95% intervals. We propose an alternative approach based on approximating the posterior with a stochastic sample from the marginal posterior. Because the the imputed paths were drawn from [***μ***|**y**], a sample from [***θ***|**y**] can be obtained by drawing samples from each [***θ***|***μ***_*r*_] and collecting them into one large sample. Then one can simply calculate a credible interval from the joint sample. This of course also applies to any function of the parameters as in any posterior sample. However, note, if different samples sizes are obtained for each [***θ***|***μ***_*r*_] then for each sample less than the largest, one can resample with replacement to obtain comparable weighting for all imputations. If GLM or GAM software is used to fit the model, one could easily approximate 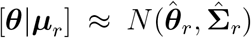, the large sample MLE or posterior distribution (see Section 4.2).

Another aspect of movement model inference using process imputation is that has received relatively little attention is that of model selection. In the first CTMC paper Hanks et al. (2015) describes use of a regularization prior (penalty) to shrink insignificant coefficients to zero, such that selection can be accomplished for nested models. However, there has been little discussion in the imputation literature with respect to selection from a discrete set of models. There have been previous proposals to perform model selection using *ad hoc* AIC weighting (Nakagawa and Freckleton, 2011). However, we propose maintaining the Bayesian paradigm inspiring the imputation approach and use posterior model probabilities to perform selection and model averaging to form a coherent process in analysis.

To begin formulating Bayesian model selection we now suppose that there is a set of *P* models, 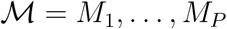 for the movement rate, 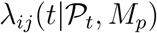 from which the likelihood, 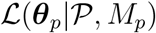, is formed. Now, using the same decomposition as (12), we can formulate

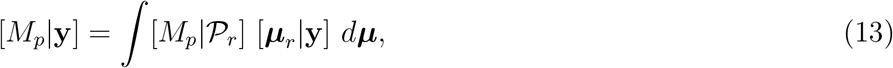

where,

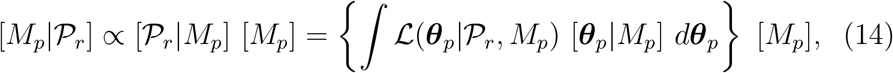

[***θ***_*p*_|*M*_*p*_] is the parameter prior distribution under model *M*_*p*_, and [*M*_*p*_] is the prior model probability. Bayesian model selection has a large volume of literature devoted to its methodology. However for application within the imputation framework, we seek a method to evaluate 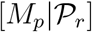 that is computationally efficient even if it is approximate, as the whole model selection procedure needs to be replicated for each ***μ***_*r*_. Therefore, we propose using Laplace approximations to equation (14; Kass and Raftery 1995),

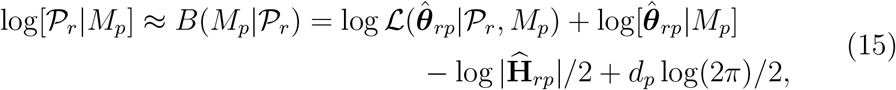

where 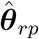 is the posterior mode of 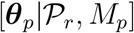, 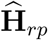 is the Hessian matrix of the negative log posterior density, and *d*_*p*_ is the dimension of ***θ***_*p*_. Kass and Raftery (1995) also note that the approximation is still of the same order if the MLE of ***θ*** is used provided the prior exerts little influence on the posterior relative to the likelihood. The benefit of that is one can use standard GLM software to fit the model without worrying about the prior distribution. However, proper priors are still necessary (highly advised) for terms that are being selected over (Bayarri et al., 2012).

Placing all the pieces back together we obtain an approximation for the posterior model probability that we can use for model selection

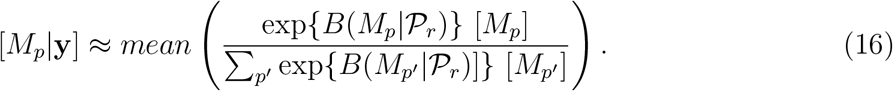

The Laplace approximation allows us to have a coherent approach to averaging an information criterion within the imputation framework.

## 4 Northern Fur Seal Pup Migration

To demonstrate analysis of movement data, we analyzed telemetry locations from 15 female northern fur seal pups tagged in the Pribilof Is., Alaska prior to their annual migration. Female ‘355’ from this data set was previously analyzed by Johnson et al. (2008) using a continuous time random walk (CTCRW). However, the previous analysis did not include any habitat related covariates to associate with movement. So, here we extend their analysis with the use of spatio-temporal covariates. Code for reproducing this analysis is available in an online appendix [^2^ 2Upon acceptance a DOI will be issued for the repository, for now, code can be accessed at: https://github.com/dsjohnson/ctsmc_movement]. These data were analyzed using the R Statistical Environment (R Core Team, 2018) due to the availability of all necessary spatial, data manipulation, movement modeling, and GLM fitting functions, however, nothing precludes development in other environments that contain spatial data manipulation and GLM capabilities.

### 4.1 A dynamic environment

Environmental covariates were defined on a hexagonal spatial grid covering the domain of the pup tracks that departed during fall 2005. For the process imputation, 20 tracks were generated using draws from the posterior distribution of locations every 15 minutes from a continuous-time correlated random walk model with a correlated random drift process (Johnson et al., 2011) using the R package `crawl` (Johnson and London, 2018) (Figure 1). Other methods of approximating the imputation distribution, [***μ***|**y**], could be used as well, e.g., basis function movement models (Hooten and Johnson, 2017; Scharf et al., 2017; Buderman et al., 2015). A regular hexagonal lattice, with 40 km center point lateral separation, and associated hexagonal polygons (“cells”) were generated. Only those cells that contain part of an imputed track or neighbors of those cells are necessary for fitting, so, only those cells are retained hereafter (see Figure 2 for an example with data from female 355). Model covariates were defined on the hexagonal grid using environmental variables obtained from remote sensing observations or atmospheric reanalysis. Along with all of the data, an animation of the spatio-temporal environmental variables is available in the online repository [^3^ https://figshare.com/account/home#/collections/3995112].

**Figure 1:**
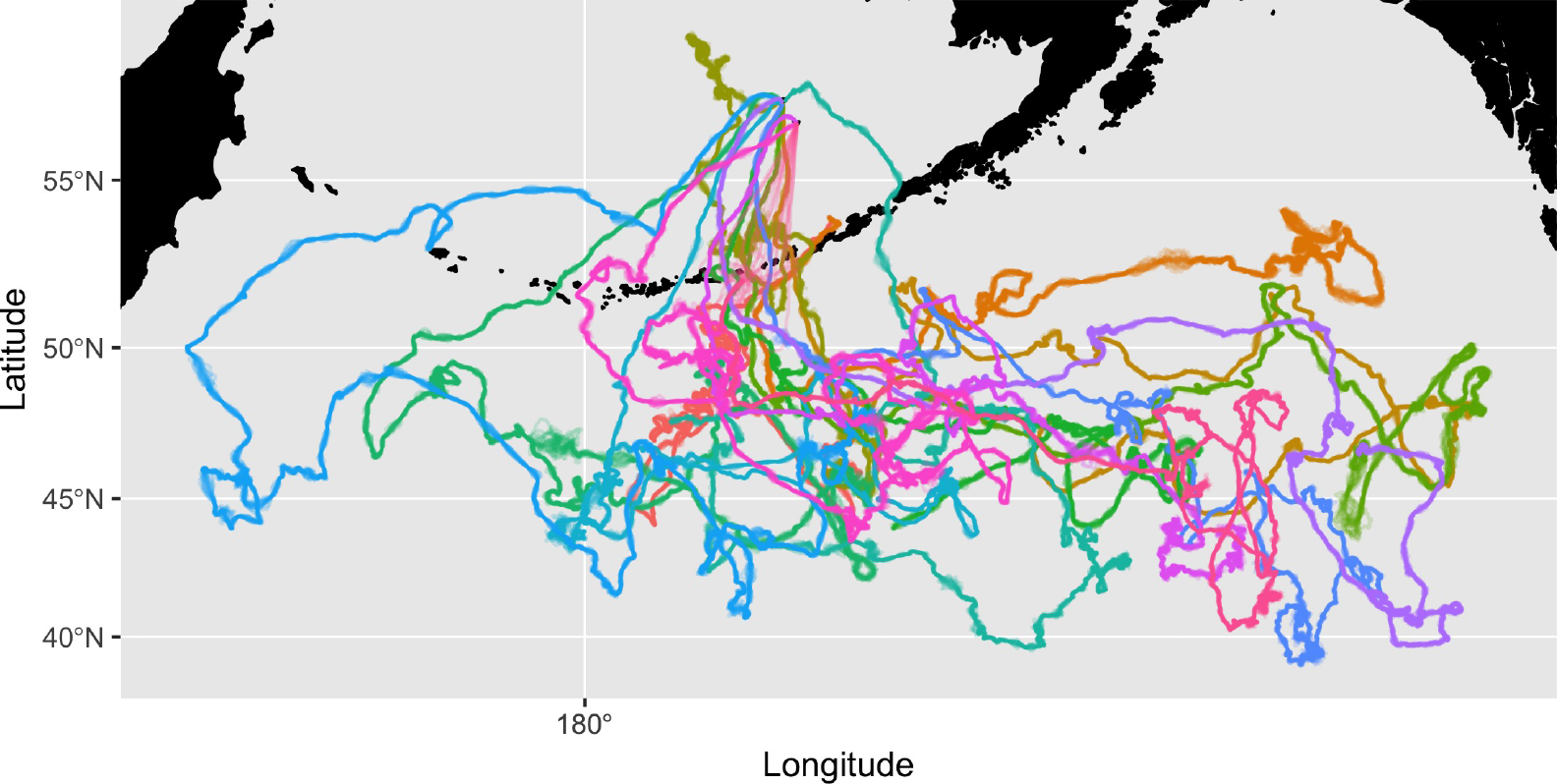
Imputed paths for analysis by the CTSMC model. Each color represents a different northern fur seal pup. Each imputation contains 20 paths for each individual.

**Figure 2:**
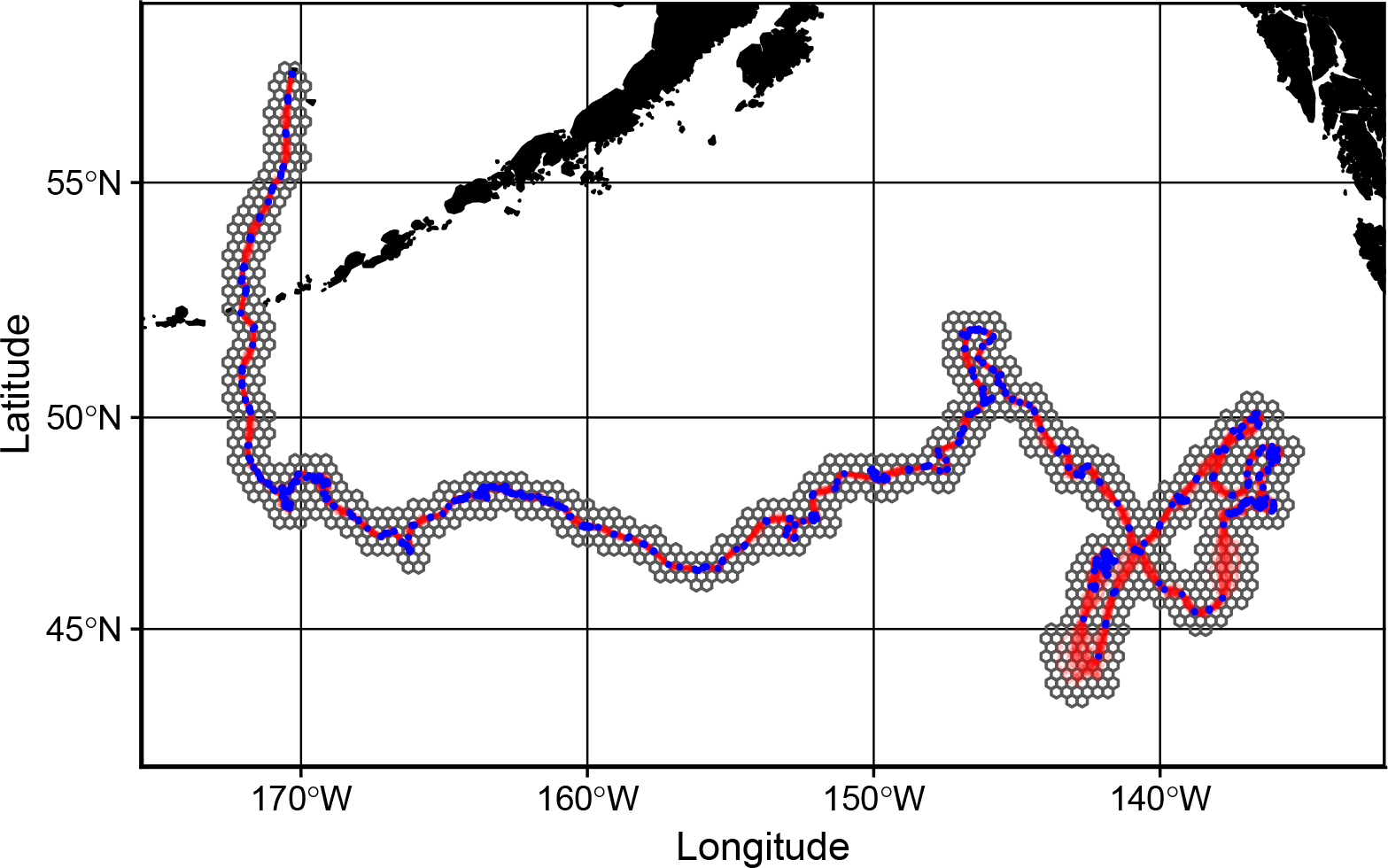
Area of deployment for the northern fur seal 355. Blue points indicate the observed locations recorded from the telemetry device. Red lines indicate the paths used for the process imputation methodology. The hex grid are the discrete locations used for CTSMC movement analysis.These hexes are all the ones used for animal 355. Not all of these are used for each imputation run.

Estimates of geostrophic ocean surface currents - slowly-varying oceanic motions (cm/s) forced by mesoscale (~10-200 km) gradients of sea surface height (Vallis, 2006) - were obtained from the AVISO All-Satellite Absolute Dynamic Topography (ADT) product (http://aviso.altimetry.fr/index.php?id=1269). The ADT product provides a smoothed, gridded estimate of the sea surface height at 1/4° spatial and daily temporal resolution based on measurements of the sea level anomaly from orbiting radar altimeters, combined with estimates of the mean sea surface obtained from *in situ* hydrographic measurements (Rio et al., 2013). At each grid point, *s*, the surface geostrophic current ***x***_*curr*_(*s*, *t*) = (*u*_*curr*_(*s*, *t*),*v*_*curr*_(*s*, *t*))*′* has a component *u*_*curr*_ in the east direction along unit vector and *v*_*curr*_ in the north direction.

Atmospheric surface (10 m height) wind estimates ***x***_*wind*_ = (*u*_*curr*_(*s*, *t*),*v*_*curr*_(*s*, *t*))*′* (in m/s) were obtained from the National Centers for Environmental Prediction/National Center for Atmospheric Research Reanalysis 1 (R1) product (http://www.esrl.noaa.gov/psd/data/gridded/data.ncep.reanalysis.html). This product gives estimates of atmospheric variables at 6-hourly resolution on a roughly 2° global spatial grid and is derived from assimilating observational data into a global forecast model (Kalnay et al., 1996). Comparison of R1 to independent observations in the North Pacific Ocean has shown that, away from topographic features, R1 winds are well-correlated with observed winds with a weak positive bias in magnitude (Ladd and Bond, 2002).

Sea surface temperature (°C) estimates, *x*_*sst*_(*s*,*t*), were obtained from the NOAA Optimal Interpolation V2 High Resolution dataset (http://www.esrl.noaa.gov/psd/data/gridded/data.noaa.oisst.v2.highres.html).

These estimates are derived from *in situ* data and the Advanced Very High Resolution Radiometer satellite instrument (Reynolds et al., 2007) and provided on a 1/4° global grid at daily temporal resolution.

Each of the environmental variables described above was spatially interpolated to the hexagonal grid cells at 6-hourly intervals. If a cell overlapped a grid boarder, the weighted average value was used based the areas contained in the hex. The 6-hour interval corresponds to the shortest resolved interval among the available variables (R1 winds); for variables obtained at daily resolution, values are repeated in multiple time intervals in each cell. For the gradient based variables, the covariates for cell transitions were defined by the projection of the gradient vector to neighboring cell centroids, that is,

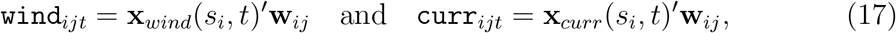

where *s*_*i*_ is the location of the *i*th hex centroid and **w**_*ij*_ is a unit length vector pointing from the hex *i* centroid to the hex j centroid (Hanks et al., 2015). SST was treated as a motility covariate to test whether pups might adjust speed if they encounter favorable conditions with respect to SST, `SST`_*it*_ = *x*_*sst*_(*s*_*i*_,*t*)

In addition, to wind and current, we defined three additional “kinematic” covariates to account for the possible effects of any background drift/movement tendency independent of environmental variables:

- 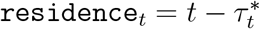
- 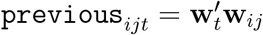,
- `north`_*ij*_ = (0, 1)*′***w**_*ij*_,
- `east`_*ij*_ = (1,0)*′***w**_*ij*_,

where 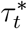 is the time of the previous transition (to the current cell), **w**_*t*_ is the unit vector of the last transition prior to *t*. `north` and `east` were included to account for pup migration tendency. The `residence` variable is the residence time within the cell (up to *t*). `previous` was added to reflect correlated movement on a finer scale than general migration patterns and is the direction of the previous cell transition. It is typical of animal movement that individuals tend to move in the same direction for some time in a correlated fashion.

In addition to the cell transition times and covariate changing times, locations on each imputed path, ***μ***_*r*_, were generated at 15 minute intervals to approximate the likelihood integral. Thus, for each quadrature time on each imputed path, *q*_*l*_, a six row data set (*j* = 1 …, 6) was created with variables: **z** = *Z*_*lj*_, `delta` = *q*_*l*_ − *q*_*l*−1_, `time` = *q*_*l*_, 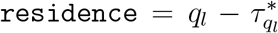, 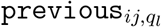, `north`_*ij*_, `east`_*ij*_, 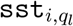, 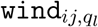, and 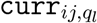. The data set was then concatenated over all *q*_*l*_ within an imputation, this forms the model data, say ‘`glm_data`’ for the *r*th imputation. From this data all models can be fitted using, say `glm` or `gam` with `family=‘poisson’`.

### 4.2 Model formulation, estimation, and selection

In pseudo-R notation, we defined the kinematic ‘Base’ movement model to be ~ `log(residence)*previous + B:north + B:east`. The B notation refers to a basis matrix used to describe the smoothly varying (over time) coefficients with which they interact. Here, the λ_0_ term in equation (7) is modeled with a `log(residence)*previous` interaction effect to account for the synergistic effect of kinetic movement on the base residence time within a cell. That is, if there is a strong relationship between the previous movement and the current one, then the residence time is likely to be longer because the animal is probably transiting the entire cell. However, if the animal is going back and forth across a border, residence time will likely be short and relationship to the previous movement will likely be negative. Because of the length of deployment, it is natural to inquire whether an individual’s movement in response to the environment changes over the course of time. The `previous` and `residence` terms were modeled as constant due to the fact that initial investigation showed no improvement by allowing their effect to vary over time. All other model terms were modeled as time varying. To accomplish this we used a radial basis function model for ***δ***(*t*) and γ(*t*), that is,

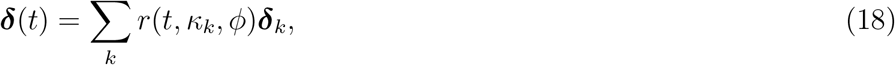

where *r*(*t*,*κ*_*k*_, *ϕ*) = exp{(*t* − *κ*_*k*_)^2^/*ϕ*^2^}, *K*_*k*_ is a knot in [0,*T*], ***δ***_*k*_ are the basis weights, and *ϕ* is a basis scaling parameter. Herein we chose 3 equally spaced interior knots plus 2 additional knots at equal spacing before and after [0, *T*] for a 5 basis model. The scalar *ϕ* set to the knot spacing interval.

For this analysis we used the R package `mgcv` and the `gam` function to estimate the posterior mode 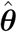, and obtain the Hessian matrix 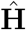, for the Laplace approximation in the posterior model probability (PMP) calculation as well as the approximating the conditional posterior

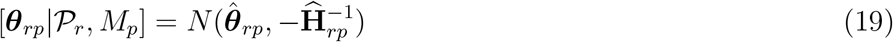

used for effect inference. To increase computational efficiency, for each animal fits were completed in parallel over the 20 imputed paths. This was accomplished with use of the R packages: `future` (Bengtsson, 2018), `doFuture` (Bengtsson, 2017), and `foreach` (Microsoft and Weston, 2017).

The posterior model probability (PMP) approximation to select between the 8 models composed of all subsets of the environmental variables (Table 1). For each model and imputation, we used the prior distribution

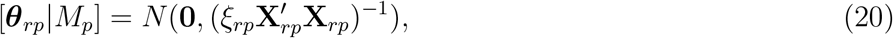

where **X**_*rp*_ is the design matrix for model (*r*,*p*) and *ξ*_*rp*_ is a scalar. A flat prior distribution was used for *ξ*_*rp*_ in each model.

**Table 1:** Model set and population average posterior model probabilities. The model column list the models in the model set considered for this analysis. The base model is, in R notation, ~ `log(residence) * previous + B:north + B:east`. The notation B indicates a radial basis matrix for allowing temporally varying coefficients. PMP column is the average PMP over all 15 animals. The range column is the range of PMP values over animals.

| Model No. | Covariates | Avg. PMP | Range |
| --- | --- | --- | --- |
| 1 | **base** | 0.003 | [0.000–0.040] |
| 2 | **base + B:wind** | 0.714 | [0.028–1.000] |
| 3 | **base + B:current** | 0.004 | [0.000–0.053] |
| 4 | **base + B:sst** |  |  |
| 5 | **base + B:wind + B:current** | 0.275 | [0.000–0.961] |
| 6 | **base + B:wind + B:sst** | 0.004 | [0.000–0.049] |
| 7 | **base + B:current + B:sst** |  |  |
| 8 | **base + B:wind + B:current + B:sst** | 0.001 | [0.000–0.011] |

### 4.3 Results

There was strong evidence for an influence of surface winds on movement with the posterior probability of the wind only model close or equal to 1 for 10 of 15 animals (Table 2). Of those animals in which the wind only model was not the maximum *a posteri* model, the model with wind and geostrophic current effects was the maximum PMP model for 3 of the remaining 5. Finally, the remaining 2 animals were basically split between models 2 and 5.

**Table 2:** Posterior model probabilities (PMP) for the CTSMC movement analysis. All models included the ‘Base’ model for movement, where `base = log residence * previous + B:north + B:east`. Model numbers correspond to rows and associated models in Table 1.

| ID | Model |  |  |  |  |  |  |  |
| --- | --- | --- | --- | --- | --- | --- | --- | --- |
|  | 1 | 2 | 3 | 4 | 5 | 6 | 7 | 8 |
| 330 |  | 0.14 |  |  | 0.86 |  |  |  |
| 335 | 0.04 | 0.34 | 0.05 |  | 0.57 |  |  |  |
| 349 |  | 1.00 |  |  |  |  |  |  |
| 351 |  | 0.92 |  |  | 0.08 |  |  |  |
| 355 |  | 0.99 |  |  | 0.01 |  |  |  |
| 362 |  | 0.99 |  |  | 0.01 |  |  |  |
| 367 |  | 0.77 |  |  | 0.23 |  |  |  |
| 377 |  | 1.00 |  |  |  |  |  |  |
| 380 |  | 0.50 |  |  | 0.45 | 0.05 |  |  |
| 381 |  | 1.00 |  |  |  |  |  |  |
| 385 |  | 1.00 |  |  |  |  |  |  |
| 388 |  | 1.00 |  |  |  |  |  |  |
| 394 |  | 1.00 |  |  |  |  |  |  |
| 405 |  | 0.03 |  |  | 0.96 |  |  | 0.01 |
| 408 |  | 0.05 |  |  | 0.95 |  |  |  |

We will now take a look at the temporally varying effects of animal ‘355’. Here we will examine the effects for model 8 even though model 2 was the highest PMP model for this animal. However, it is instructive to see the general pattern of all covariates (Figure 3). For animal 355, the wind effect is overall positive throughout the course of the deployment implying that the animal generally travels with the prevailing winds although, the strength of this effect seams to vary with time. There also seems to be a generally positive effect of geostrophic current on the rate of movement, although for only a small fraction of the deployment is it significantly positive according to the 95% credible interval. This small interval is not enough to make the covariate overall significant as the wind only model has PMP = 0.99. SST has a negative but insignificant effect meaning that the animal tends to slow in warmer seas, but it is highly uncertain. The kinetic nuisance effects, north and east are also illustrated and describe the general trend in movement as expected, first southerly movement followed by generally easterly movement.

**Figure 3:**
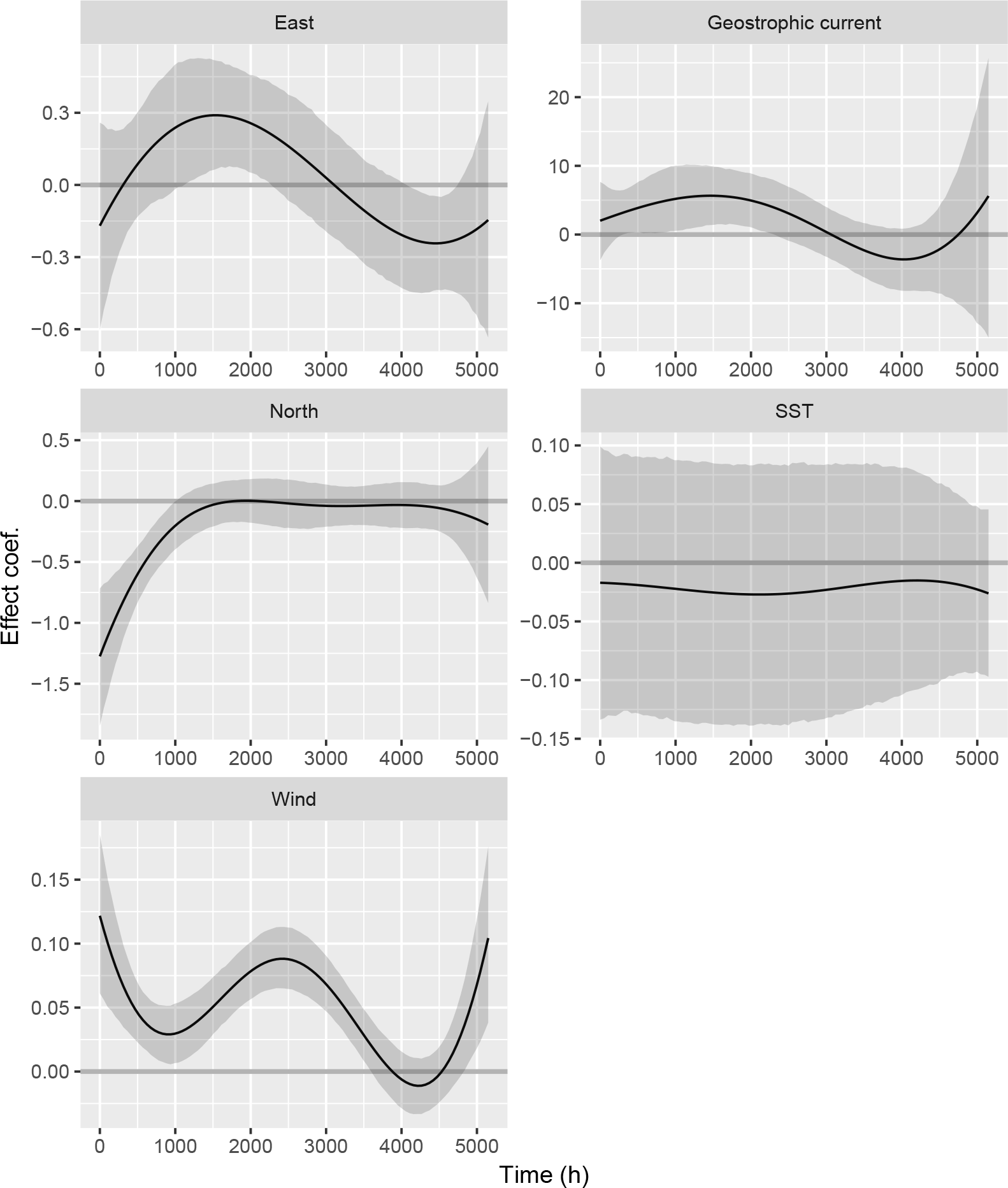
Time varying coefficient estimates for the CTSMC model fit to individual 355. Estimates and credible intervals were calculated using the process imputation methodology presented in Section 3.3 using a normal approximation to the posterior for each imputation.

## 5 Discussion

Here we’ve presented an extension to the CTMC model of Hanks et al. (2015) for animal movement modeling. In addition to the ability to handle dynamic habitat covariates the semi-Markov version (CTSMC) relaxes the assumption of exponentially distributed residence times for each cell. However, even after the extensions, the Poisson likelihood approximation still holds with only slight considerations, that is at quadrature points without cell transition, none of the *Z*_*lj*_ values are set to 1. It can be thought of as the absence of movement at that time. Thus, there is no additional difficulty in practice to fit a temporally dynamic model (covariates and coefficients) as it is to fit the time-constant habitat CTMC model.

To fully account for path uncertainty in the analysis, we used a very similar 2-stage imputation approach used by Hanks et al. (2015). However, an alternative is to use a ‘stacked’ likelihood (e.g., White et al. 2011; Hanks and Hughes 2016). Thus, one can make inference using this weighted likelihood and avoid having to recombine separate model inference. However, the spatio-temporal data sets built using the methodology of Section 4.1 might often preclude this as the complete data set might become too large to deal with efficiently for, say, a complex GAM model. It would be more efficient to analyze each imputed data set separately and recombine the results in a buffered fashion. In this way the separate fits could can be parallelized for increased computational speed as we have done in the example herein.

In addition to extension of the CTMC model to a semi-Markov version, we have also explored model selection in the context of a 2-stage imputation approach. To our knowledge, this general topic has not been explored outside of Nakagawa and Freckleton (2011), and only with respect to AIC model averaging therein. Here we propose a fully Bayesian model section approach in an imputation setting using an approximated version of the posterior model probability (PMP). PMPs can then be averaged over imputations in accordance with the Bayesian interpretation of multiple imputation to account for path uncertainty in selection.

The Laplace approximation used in the PMP calculations allows for computational efficiency over all of the imputed paths. One might question whether the approximation provides values close enough to the true marginal distribution 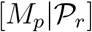. While there is no definitive way to know whether it is the case for any particular data set, Kass and Raftery (1995) note that sample sizes < 5*d*_*p*_ are worrisome, while those > 20*d*_*p*_ should be sufficiently accurate. The question for these models is, “what is the sample size?” Volinsky et al. (1997) notes that the appropriate quantity for proportional hazard models is the number of observed events in the data. In the CTSMC models this would be the number of observed cell transitions *n*_*r*_ = ∑_*l*_ *Z*_*lj*_ for each 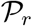. In the fur seal example the largest model (model 8) contains 30 parameters, thus sample sizes > 600 should cause little worry for any of the models fitted here. For the fur seal data, only 1 animal fell below this threshold and its minimum was *n*_*r*_ = 560 over all 20 imputed paths. So, we believe that *n*_*r*_ was sufficient in all animals for this analysis. However, this is something to keep in mind for other data. It should be noted that for each imputation full PMPs must be calculated so that imputation dependent normalization constants in (14) cancel. If one uses, say, *B* (eqn. 15) or BIC, alone the value will not be comparable over imputations because 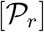 has not been included. This is essentially the same reason that comparing AIC or BIC values between datasets is not valid.

The CTSMC model presents a computationally feasible approach for making inference with respect to the influence of dynamically changing habitat on animal movement. The ratio λ_*ij*_/Λ_*i*_ essentially represents a discrete choice resource selection model (McCracken et al., 1998) with the additional inference on rate of movement modeled as well. The imputation methodology allows one to disentangle the selection from the rate of movement by regularizing the modeled locations of the animal. Thus, we recommend the CTSMC approach as a general purpose resource selection and movement analysis tool.

## Acknowledgments

The findings and conclusions in the paper are those of the authors and do not necessarily represent the views of the National Marine Fisheries Service, NOAA. Reference to trade names does not imply endorsement by the National Marine Fisheries Service, NOAA.

